# AI-guided design and *ex vivo* validation of nanobodies targeting aggregation motifs of intrinsically disordered protein tau

**DOI:** 10.64898/2026.04.01.715983

**Authors:** Binita Rajbanshi, Anuj Guruacharya

## Abstract

Intrinsically disordered proteins (IDPs) represent major yet challenging therapeutic targets in neurodegenerative disease due to their conformational heterogeneity and aggregation-prone behavior. Tau protein is a prototypical IDP that forms pathological aggregates in Alzheimer’s disease and related tauopathies. Despite extensive clinical efforts, tau-directed monoclonal antibodies have demonstrated limited efficacy. Concurrently, single-domain antibodies (nanobodies) have been gaining importance because of their small size and membrane penetrating capabilities. New design paradigms are therefore required for nanobodies to enable precise targeting of disease-relevant conformations. Here, we describe a biophysical modelling and AI-guided nanobody discovery targeting the VQIVYK motif of tau, which constitutes the structural core of neurofibrillary tangles in Alzheimer’s Disease. Biophysical modelling-based target analysis identified low-energy conformational states of VQIVYK. These conformational insights were used to guide AI-driven nanobody design of CDR3 loops. Starting from a nanobody scaffold, we generated 145 candidate nanobodies through systematic backbone sampling and neural network-guided sequence design, followed by multi-dimensional computational prioritization. Two candidates demonstrated robust binding to synthetic full tau protein in ELISA binding assays, achieving binding indices of 148.9% and 140%, relative to reference controls. Notably, one candidate also exhibited strong reactivity in post-mortem Alzheimer’s disease human brain tissue, with a binding index of 236.1%, exceeding that of the positive control (222.9%). Structural analysis indicates that our nanobodies’ engineered CDR3 engages VQIVYK through optimized aromatic and hydrophobic interactions. Together, these findings establish a proof-of-concept for biophysics-guided, AI-guided nanobody engineering against IDPs and identifies them as a promising lead for tau-targeted single domain antibody development.

## Main

Intrinsically disordered proteins (IDPs) constitute a major class of disease-relevant targets that have historically resisted conventional therapeutic strategies [1,2]. Unlike globular proteins with stable tertiary structures, IDPs populate dynamic conformational ensembles and undergo pathological self-assembly into aggregated states [3–5]. In neurodegenerative diseases, such aggregation underlies protein misfolding pathologies, yet the transient and heterogeneous nature of IDPs poses fundamental challenges for therapeutic targeting [6–8]. Traditional antibody discovery approaches, which rely largely on empirical screening against static epitopes, are poorly suited [9–11] to engage the conformationally diverse and aggregation-prone species that drive disease progression.

Tau protein exemplifies these challenges. As an intrinsically disordered microtubule-associated protein, tau adopts multiple conformational states and aggregates into paired helical filaments and neurofibrillary tangles, defining pathological features of Alzheimer’s disease and related tauopathies [12,13]. Despite extensive clinical development efforts [14–17], tau-directed antibodies have thus far failed to produce disease-modifying benefits . Key limitations include insufficient engagement of pathogenic tau species within the central nervous system, driven by limited brain exposure [18,19], inadequate selectivity for disease-relevant conformations [20,21], and poor accessibility of aggregation-prone epitopes that become structurally buried within fibrillar assemblies [22,23]. These shortcomings underscore the need for alternative antibody formats and discovery strategies capable of targeting pathological tau with greater precision.

At the molecular level, the VQIVYK motif of the tau protein which is a hexapeptide sequence (Valine - Glutamine - Isoleucine - Valine - Tyrosine - Lysine) corresponds to amino acid residues 306-311 in the longest canonical isoform of tau (2N4R, UniProt P10636-8) [24,25]. This region lies at the beginning of the third microtubule-binding repeat domain (R3) and constitutes the core of the aggregation-prone region of tau that nucleates β-sheet formation [12,26]. The VQIVYK sequence has been demonstrated in numerous structural and biochemical studies to play a central role in the formation of paired helical filaments (PHFs) and straight filaments (SFs), which together comprise the neurofibrillary tangles characteristic of Alzheimer’s disease and related tauopathies [27,28].

Cryo-electron microscopy (cryo-EM) structures of *ex vivo* tau fibrils consistently identify the VQIVYK region as being at the interface of the β-sheet-rich fibril core [29–31]. Structures solved from Alzheimer’s disease, progressive supranuclear palsy, corticobasal degeneration, and chronic traumatic encephalopathy brain tissue all retain the VQIVYK motif in a flat, extended β-strand geometry at the steric zipper interface, underscoring its conservation across diverse tauopathies [27,31,32]. Peptides corresponding to VQIVYK alone are sufficient to induce aggregation *in vitro* and are capable of seeding intracellular tau aggregation in biosensor cell lines [33,34]. Furthermore, mutations or post-translational modifications near this motif alter aggregation propensity, further highlighting its critical role [35,36].

Given its central function in disease-specific aggregation and limited exposure in physiological tau, the VQIVYK motif represents an ideal epitope for selective targeting of pathological tau conformers [12,16,27,37]. However, its hydrophobic nature and location within the fibril core pose challenges for conventional antibody discovery platforms [38,39].

Single-domain camelid antibodies (VHHs, or nanobodies) are particularly well suited for targeting structurally constrained epitopes [40–42]. Their small size (∼15 kDa), high thermostability, and extended CDR3 loops enable access to epitopes that may be sterically occluded to conventional immunoglobulins, including those embedded within aggregated protein assemblies [43–46]. Nanobodies can be expressed at high yield in microbial systems, are amenable to genetic fusion and multimerization, and have demonstrated blood–brain barrier penetration in preclinical models, properties that collectively position them as an attractive modality for precision targeting of pathological tau. Notably, the WIW nanobody (PDB: 8FQ7) demonstrated that a Trp–Ile–Trp aromatic triad within the CDR3 loop can directly cap the VQIVYK steric zipper interface and inhibit tau fibril seeding, establishing structural precedent for CDR3-mediated engagement of this epitope [47].

In this study, we circumvent these limitations by using in silico structural modeling and targeted design of nanobody CDR3 loops to interact with the unique shape and chemical environment of intrinsically disordered regions like the VQIVYK region of tau protein. To address the conformational heterogeneity inherent to this IDP-derived epitope, molecular dynamics simulation was used to sample 100 conformers of the VQIVYK hexapeptide, providing an ensemble representation of the target for structure-based design. Multiple structural hypotheses were tested via docking against VQIVYK-containing tau fibril models, optimizing for interface complementarity, binding energy, and CDR3 engagement. Accordingly, all nanobody sequences are determined either directly modeled to bind the VQIVYK hexapeptide motif or are designed to bind structural epitopes containing the sequence in the context of aggregated or misfolded tau.

Here, we show that the design and validation of single-domain antibodies targeting a therapeutically important intrinsically disordered region, VQIVYK aggregation motif of tau protein. By combining conformational analysis of the tau VQIVYK motif with AI-guided protein engineering to develop precision binders against pathological full-length tau. Starting from the validated WIW tau-binding nanobody scaffold [47], we employed AlphaFold2 [48] for structural modeling and ProteinMPNN [49] for generative sequence design to systematically explore CDR3 conformational space. Candidate nanobody-VQIVYK complexes were refined and prioritized using HADDOCK 2.4 [50] molecular docking, and ranked by a composite metric incorporating binding complementarity, structural stability, and developability. Lead nanobodies were evaluated for full-length tau binding *in vitro* and for recognition of pathological full-length tau species in post-mortem Alzheimer’s disease human brain tissue. This work establishes a rational, computationally driven strategy for antibody engineering against intrinsically disordered protein targets, demonstrates its application to a central aggregation motif in tau pathology, and validates the design outputs as promising leads for pathological tau-targeted single-domain antibody development.

## Results

### Target exploration of VQIVYK motif of tau

Tau is an IDP and lacks a fixed tertiary structure, so we sampled 100 conformers of the VQIVYK hexapeptide by molecular dynamics (MD) simulation. Analysis of 100 conformers of the tau VQIVYK hexapeptide revealed substantial conformational heterogeneity across the MD trajectory. Total MD energies spanned 653.15–809.62 kJ/mol (mean 727.94 ± 26.01 kJ/mol), while intramolecular energies averaged 1647.07 ± 41.91 kJ/mol, consistent with a broad and flexible conformational ensemble. Energy decomposition indicated that Coulombic 1-4 interactions were the dominant contributors to intramolecular energy (62.6%; 1030.47 ± 25.67 kJ/mol), followed by angle bending and proper dihedral terms. Bond stretching exhibited the greatest relative variability (coefficient of variation, 17.8%), reflecting sensitivity of local geometry across conformers.

Single-point RHF calculations performed on the full 114-atom peptide yielded energies ranging from -2476.044 to -2475.824 Hartree (-1,553,739.95 to -1,553,602.13 kcal/mol), corresponding to a conformational energy spread of 137.82 kcal/mol. These results indicate substantial electronic energetic diversity across aggregation-relevant conformations. Active space analysis of a reduced 20-atom fragment centered on the tyrosine aromatic system showed an even broader energetic range (181.07 kcal/mol), with mean fragment energies of −235,381.85 ± 23.21 kcal/mol, highlighting local electronic flexibility within the VQIVYK motif.

Correlation analysis revealed a modest but statistically significant positive relationship between full-molecule RHF energies and MD intramolecular energies (Pearson r = 0.31, p = 1.67 × 10^−3^; Spearman ρ = 0.32, p = 1.15 × 10^−3^), indicating partial correspondence between classical and electronic descriptions of global conformational energetics. In contrast, active space fragment energies showed no significant correlation with either full-molecule RHF or MD energies, demonstrating that local aromatic electronic structure is largely decoupled from global peptide conformation.

Notably, the lowest-energy conformers were distributed throughout the trajectory rather than temporally clustered, consistent with a heterogeneous ensemble of aggregation-competent states. A representative summary of the computational analysis is shown in **Fig. 1**.

**Fig. 1:**
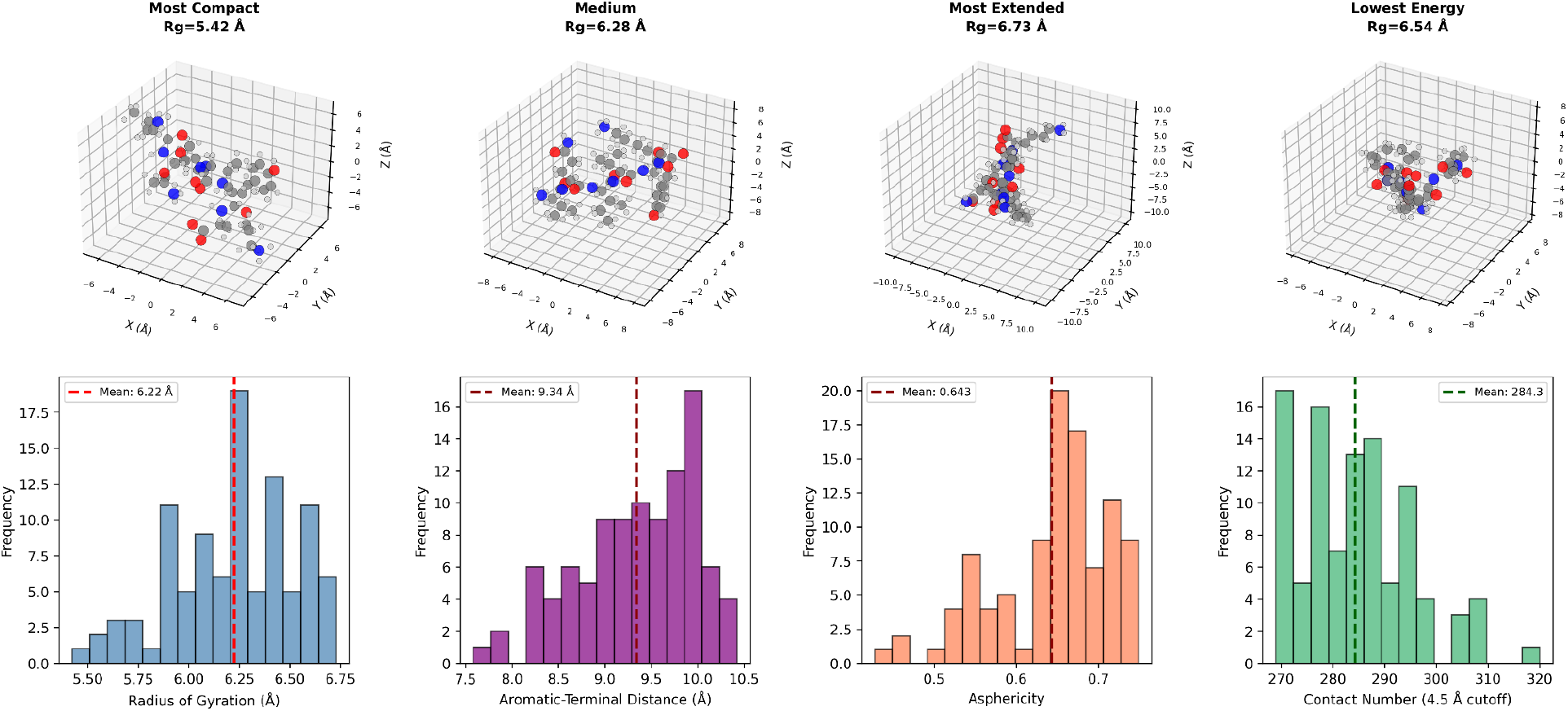
Conformational ensemble and structural characterization of VQIVYK tau peptide. Representative conformers from molecular dynamics simulations showing structural diversity across the ensemble. Distribution of key structural descriptors across 100 conformers.

### Nanobody hit screening targeting the VQIVYK epitope

Starting from the tau VQIVYK steric zipper structure (PDB 5K7N, 1.1 Å microED) and the WIW nanobody scaffold (PDB 8FQ7, 1.4 Å), we established an AI-guided design and prioritization workflow to enrich tau-binding nanobody candidates targeting this disease-relevant epitope. The VHH scaffold was held constant across framework regions (FR1–FR4) and CDR1/CDR2 loops, while sequence diversity was introduced within the CDR3 region (residues 103–109), which dominates antigen recognition. Ser102 was retained as a fixed anchor to preserve framework connectivity. ProteinMPNN was used to generate 145 candidate CDR3 sequences, restricting variation to the 7 CDR3 positions while holding the remainder of the scaffold fixed. AlphaFold2 was then used to generate three-dimensional structural models of the candidate nanobodies, enabling assessment of folding plausibility and CDR3 conformational diversity prior to docking.

AlphaFold2-predicted nanobody structures were docked to the VQIVYK-containing tau fibril using HADDOCK 2.4 rigid-body docking followed by semi-flexible refinement and energy minimization in explicit water. Docking analyses identified consistent binding poses in which the CDR3 loop engages the VQIVYK motif through a combination of hydrogen bonding and hydrophobic interactions **(Fig. 2A)**. Each complex was evaluated with a weighted composite metric comprising binding score (40%), stability score (30%), and developability score (30%). Across the full library, the composite score followed a roughly normal distribution with a mean of 0.634 ± 0.051, ranging from 0.509 to 0.809 **(Fig. 2B)**. The binding score, which assessed position-specific complementarity to the VQIVYK motif, aromatic content for π-stacking with Tyr(305), and electrostatic complementarity with Lys(311), exhibited the widest dynamic range among the three scoring components and served as the primary discriminator between high- and low-ranking candidates.

**Fig. 2:**
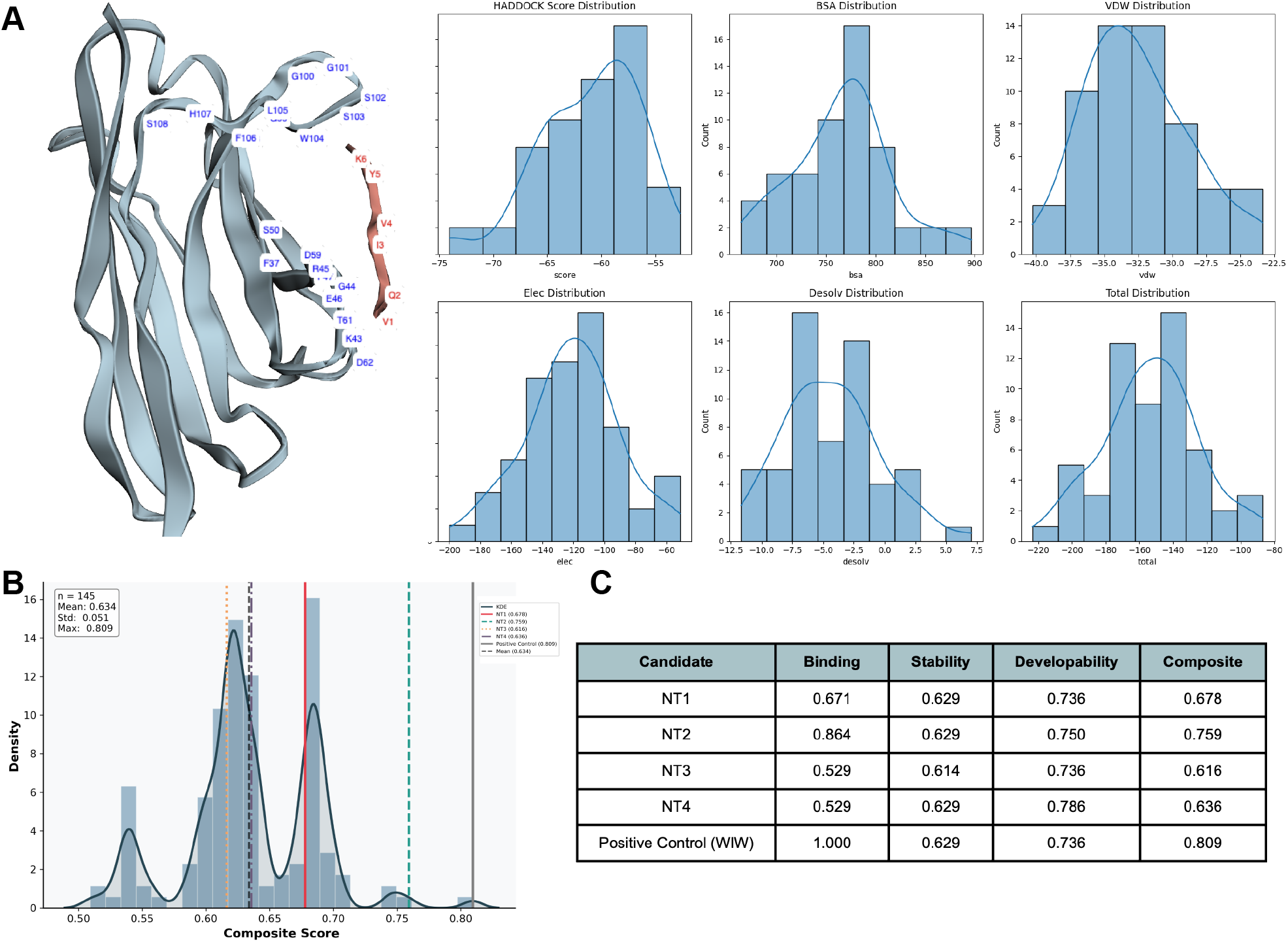
Computational design and selection of VQIVYK-targeting nanobodies. **(A)** A representative docked complex generated using HADDOCK shows the nanobody bound to the VQIVYK region of tau. A distribution of predicted binding energies (kcal/mol) for top-ranked nanobody candidates. **(B)** Scoring comparison showing NT2 achieved the highest composite score (0.759) among selected candidates and positive control scaffold. **(C)** Composite score distribution across 145 candidates. Selected candidates (dashed lines) span the upper quartile.

From this AI-guided screening process, four representative nanobody candidates were selected for initial experimental evaluation. NT1 scored a composite of 0.678. NT2 achieved the highest composite score among the four designed candidates (0.759) and the highest binding score (0.864). NT3 was selected for its CDR3 sequence divergence from NT1 and NT2 (composite of 0.616). NT4 scored a composite of 0.636 and achieved the highest developability score (0.786) among the four candidates, with no predicted deamidation motifs, glycosylation sequon, or unpaired cysteines **(Fig. 2C)**.

The parental WIW nanobody CDR3 served as the positive control and scored a composite of 0.809, the highest in the library. The Trp104-Ile105-Trp106 aromatic triad that mediates steric zipper capping was partially conserved across all four designed candidates through retention of Trp104. NT1 and NT2 further recapitulated the hydrophobic character of Ile105 with Leu105, while NT3 retained Ile105 directly. NT4 adopted Val105 as a compact hydrophobic alternative. NT2 most closely approached the positive control in binding score (0.864 vs 1.000), driven by a near-complete match of aromatic and electrostatic features at key CDR3 positions. NT4 exceeded the parental developability score (0.786 vs 0.736), while NT2 also improved upon it (0.750 vs 0.736). The four selected candidates sampled distinct CDR3 chemistries. Aromatic-hydrophobic (NT1), aromatic-electrostatic (NT2), polar-aromatic (NT3), and charged-hydrophobic (NT4) providing broad coverage of the sequence space compatible with VQIVYK binding for subsequent experimental validation.

### *In-vitro* and *ex-vivo* neuropathological binding validation

Nanobodies were recombinantly expressed and subjected to rigorous *in-vitro* binding validation using enzyme-linked immunosorbent assays (ELISA). To establish specificity and benchmark performance, candidates were evaluated alongside a VQIVYK-binding single domain antibody (positive control, PDB: 8FQ7),and a non-tau targeting nanobody (negative control). All lead candidates demonstrated selective, concentration-dependent binding to tau protein, confirming successful computational design translation to functional biomolecules. Binding performance was quantified using a normalized binding index, with a commercially available tau antibody serving as the reference standard (100%). This normalization approach enabled direct comparison across multiple independent experiments and facilitated objective candidate ranking. The negative control nanobody yielded minimal signal (2.4%), confirming assay specificity and establishing a robust signal-to-noise ratio suitable for discriminating true binders from non-specific interactions.

Among the tested candidates, NT1 and NT2 nanobodies emerged as superior binders, achieving normalized binding indices of 148.9% and 140%, respectively substantially exceeding the reference antibody standard **(Fig. 3A)**. These candidates demonstrated consistent performance across biological replicates, indicating reproducible tau recognition. Additional variants NT3 and NT4 exhibited intermediate binding profiles (76% and 53.3%, respectively). The broad dynamic range observed across candidates validated ELISA as an effective primary screening platform for this discovery campaign. The differential binding performance observed across the nanobody panel, coupled with negligible background signal from negative controls, confirmed both the specificity of tau engagement and the discriminatory power of the screening assay. These results validate the power of computationally designed nanobodies as a complementary approach to conventional antibody discovery platforms for targeting structurally constrained epitopes within IDPs. Further these results provided high confidence for prioritizing NT1 and NT2 as lead candidates for subsequent hit-to-lead optimization, including affinity maturation, epitope mapping, and functional characterization in cellular tau aggregation models.

**Fig. 3:**
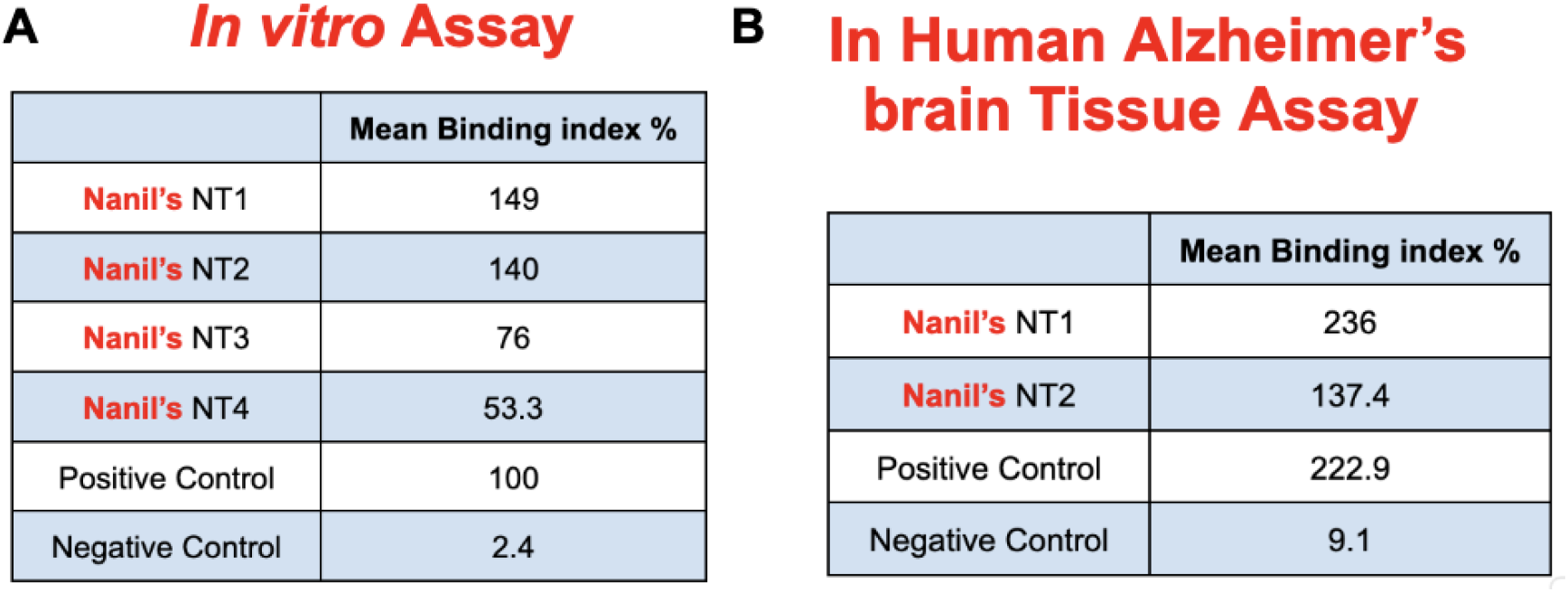
*In vitro* and *ex vivo* binding of nanobody drug candidates in post-mortem human Alzheimer’s disease brain tissue. **(A)** Normalized binding index (%) measured by ELISA for Nanil’s tau-targeting VHH candidates (NT1-NT3) relative to a publicly available tau antibody used as a positive reference control (set to 100%), and a negative control. **(B)** Normalized binding index (%) measured in human brain tissue sections for Nanil’s tau-targeting VHH candidates, a positive reference control, and a negative control.

To assess disease relevance in a native pathological context, binding was further evaluated in post-mortem human Alzheimer’s disease brain tissue **(Fig. 3B)**. Using the same normalized binding index, Nanil’s NT1 demonstrated robust tissue binding (236.1%), exceeding that of the positive reference control (222.9%), while Nanil’s NT2 nanobody showed moderate binding (137.4%). In contrast, the negative control exhibited minimal signal (9.1%), indicating low nonspecific background. Replicate measurements revealed consistent binding index distributions for Nanil’s NT1 and NT2 nanobodies, supporting reproducibility across tissue samples. Collectively, these data demonstrate that the lead mAb candidates, particularly NT1, recognize disease-relevant tau species in human brain tissue, providing strong translational support for advancement into hit-to-lead optimization, including affinity maturation, developability assessment, and downstream functional evaluation to de-risk progression toward *in vivo* studies.

## Discussion

This study for the first time demonstrates that biophysical modeling and computationally driven nanobody engineering can overcome long-standing barriers in targeting IDPs such as tau, whose VQIVYK aggregation motif has remained largely inaccessible to conventional antibody discovery platforms due to its conformational heterogeneity and burial within the fibril core. The repeated failure of tau-directed immunotherapies in clinical trials [14–17] has underscored fundamental limitations of epitope-agnostic approaches, namely their inability to discriminate pathological tau conformers from physiological species [14,22]. By contrast, the CDR3 design strategy employed here directly addresses this challenge, leveraging the VQIVYK steric zipper as a conformation-specific epitope to engineer nanobodies with defined binding mechanisms against pathological tau assemblies implicated in Alzheimer’s disease and related tauopathies.

Molecular modelling of the VQIVYK motif revealed substantial conformational and energetic heterogeneity, consistent with its central role in nucleating tau aggregation. The modest but significant correlation between classical MD and full-molecule electronic RHF energies indicates partial correspondence between conformational sampling and electronic stabilization, while the lack of correlation between local aromatic fragment energetics and global peptide energies highlights the decoupling of local electronic features from overall backbone conformation. These findings support a design strategy in which nanobody paratopes are optimized to engage conserved local features of aggregation-prone motifs across a heterogeneous conformational ensemble, rather than a single static structure.

Leveraging these insights, we combined AlphaFold structural modeling with HADDOCK molecular docking to systematically explore the nanobody CDR3 design space. Starting from the validated WIW tau-binding scaffold, ProteinMPNN was used to generate candidate CDR3 sequences, whose predicted structures were docked against the VQIVYK fibril to evaluate binding pose compatibility. This approach enabled rapid generation and prioritization of structurally diverse candidates while explicitly balancing binding complementarity, structural stability, and developability. Importantly, diversity-weighted selection proved essential for capturing high-performing candidates that were not strictly top-ranked by composite score, emphasizing that exploration remains critical even within rational design pipelines.

Further, experimental validation in *invitro* and Alzheimer’s disease human brain tissue confirmed the effectiveness of this strategy. NT1 and NT2 demonstrated robust binding to synthetic tau, and NT1 showed particularly strong recognition of pathological tau species in post-mortem Alzheimer’s disease brain tissue, exceeding the performance of a positive control antibody. Notably, this level of disease-relevant binding was achieved without experimental affinity maturation, underscoring the power of structure-guided, AI-enabled design for nanobody engineering. Overall, demonstrating that fully AI-designed nanobodies generated without predefined inhibitor sequences can achieve comparable or superior binding to pathological tau assemblies. This suggests that the design space for aggregation-targeting antibodies is broader than previously appreciated and can be efficiently navigated using generative computational approaches.

Several limitations warrant consideration. Binding alone does not guarantee therapeutic efficacy, and future studies will need to evaluate the effects of NT1 and NT2 on tau aggregation, seeding, and cellular toxicity, as well as assess developability parameters such as stability and immunogenicity. Nonetheless, the identification of a computationally designed nanobody with strong disease-relevant binding establishes a compelling starting point for downstream optimization.

In conclusion, this work provides the foundation and is first of its kind that integrating classical computing-informed target analysis with AI-guided nanobody design enables precise targeting of intrinsically disordered, aggregation-prone proteins. Beyond tau, this approach is broadly applicable to other neurodegenerative disease targets that have resisted conventional antibody discovery, offering a rational path toward next-generation biologics for protein misfolding disorders.

## Author contributions

BR and AG conceived and designed the study. Both authors contributed to revisions across all sections and approved the final manuscript.

